# Whole chromosome loss in tetraploid cells confers tumorigenic potential in a mouse allograft model

**DOI:** 10.1101/172668

**Authors:** Rozario Thomas, Daniel H Marks, Yvette Chin, Robert Benezra

## Abstract

Whole chromosome gains or losses (aneuploidy) are a hallmark of ~70% of human tumors. Modeling the consequences of aneuploidy has relied on perturbing spindle assembly checkpoint (SAC) components but interpretations of these experiments are clouded by the multiple functions of these proteins. Here we used a Cre recombinase-mediated chromosome loss strategy to individually delete mouse chromosomes 9, 10, 12 or 14 in tetraploid immortalized murine embryonic fibroblasts. While the aneuploid cells generally display a growth disadvantage *in vitro*, they grow significantly better in low adherence sphere-forming conditions and 3 of the 4 lines are transformed *in vivo*, forming large and invasive tumors in immunocompromised mice. The aneuploid cells display increased chromosomal instability and DNA damage, a mutator phenotype associated with tumorigenesis *in vivo.* Thus, these studies demonstrate a causative role for whole chromosome loss in tumorigenesis and may shed light on the early consequences of aneuploidy in mammalian cells.

## Introduction

While aneuploidy is associated with a high percentage of human cancers, whether it can directly promote tumorigenesis is still not firmly established. Most current models of aneuploidy rely on altering the levels of various SAC proteins, which ensure sister chromatid segregation occurs only after chromosomes are properly bi-oriented during metaphase. Several mouse models of aneuploidy have been generated that harbor hypomorphic alleles or heterozygous knockouts for key SAC proteins such as Mad2, Bub3, and BubR1 with varying tumor penetrance, spectrum and latency (Schvartzman et al., 2010; Simon et al., 2015). However, genetic loss of these SAC components is not usually observed in human tumors. In contrast, SAC components are frequently overexpressed. For instance, Mad2, a key SAC protein is overexpressed in a wide array of human tumors, and in transgenic mouse models overexpression of Mad2 can initiate and promote tumorigenesis (Sotillo et al., 2007). A major caveat of these mouse models is that SAC proteins have other non-mitotic functions such as in nuclear trafficking, transcriptional repression, apoptosis and the DNA damage response (Schvartzman et al., 2010). Therefore it is unclear whether the pro-tumor phenotype observed is a direct result of the aneuploidy generated.

Other groups have generated yeast, mouse and human cells with single chromosome gains. Haploid yeast strains harboring extra chromosomes exhibit a growth disadvantage compared to the controls and a transcriptional response similar to the yeast environmental stress response (Torres et al., 2007). Also, the growth inhibitory phenotypes of these disomic strains were due to the expression of the genes present on the extra chromosome and on the stoichiometric imbalance resulting from the supernumerary proteins (Torres et al., 2007). The aneuploid cells adapt by buffering the levels of these extra proteins and are therefore more sensitive to additional proteotoxic stress, like inhibitors of protein folding and protein degradation (Oromendia et al., 2012; Torres et al., 2007). In addition, these aneuploid strains were also found to lose other chromosomes at a high frequency and to exist in a generally unstable karyotypic state (Sheltzer et al., 2011).

Experiments using trisomic mouse embryonic fibroblasts have shown that they also have growth impairment relative to the euploid MEFs and a delay in the rate to spontaneously immortalize (Williams et al., 2008). The growth impairment has been attributed to the proteotoxic stress associated with carrying the extra chromosomes. Similar to the yeast model, these trisomic MEFs have been shown to be sensitive to proteotoxic stressors since their protein folding and degradation machinery is already under duress (Tang et al., 2011). Also, in reconstitution experiments into lethally irradiated mouse recipients, these trisomic hematopoietic stem cells (HSCs) displayed reduced fitness in comparison to the wild type HSCs (Pfau et al., 2016). The fitness and neoplastic defects in the trisomic MEFs are not overcome even with the addition of oncogenic mutations (Sheltzer et al., 2017). Recently, trisomic mouse embryonic stem (ES) cells were generated by a series of steps involving random integration of a dual drug selection cassette followed by expansion in culture to allow for an increase in copy number and finally Cre recombination to generate dual drug resistant ES cells, trisomic for a particular chromosome. These trisomic ES cells were found to be less differentiated and have enhanced teratoma forming potential in comparison to the wildtype ES cells (Zhang et al., 2016).

Human cells harboring extra chromosome(s) have a similar growth defect and exhibit a uniform aneuploidy response signature (specific pathways being up-regulated or down-regulated irrespective of the identity of the chromosome gained) (Stingele et al., 2012). Interestingly, while the DNA and mRNA copy numbers are increased corresponding to the specific chromosome gained, the protein levels, especially those that are subunits of multi molecular complexes, are maintained at normal levels. One way these trisomic or tetrasomic aneuploid cells achieve this protein homeostasis is by increasing the function of the autophagy pathway (Stingele et al., 2012). Also, human cells with extra chromosomes were shown to have replicative defects because of down-regulation of the helicase, MCM2-7 and importantly, these aneuploid cells were found to be genomically unstable (Passerini et al., 2016). Similar experiments on trisomic colorectal cancer cells also show that these aneuploid cells mis-segregate chromosomes at a higher rate than the control diploids (Nicholson et al., 2015).

While all of the studies mentioned above focus on gain of chromosome aneuploidies, to our knowledge, no groups have studied the effects of chromosome-loss mediated aneuploidy on cellular fitness and tumorigenesis. Furthermore, a comprehensive study of human tumor karyotypes revealed a strong disposition for chromosome loss rather than gains (Duijf et al., 2013). Here, we show that the loss of individual chromosomes in a tetraploid background induces further genetic instability and drives tumorigenesis in a mouse allograft model.

## Results

### Generation of Cre recombinase-mediated whole chromosome loss

In order to model chromosome loss events, we generated SV40 Large T-antigen immortalized mouse embryonic fibroblasts (MEFs) bearing a pair of inverted loxP sites (iLoxP) on one homolog of chromosome 9, 10, 12 or 14 (Ch9, Ch10, Ch12 or Ch14) (Figure 1A). As reported in other studies, these MEFs become tetraploid as a result of the Large T-antigen immortalization procedure (Hein et al., 2009; Lionnet et al., 2011) and thus the initial ploidy prior to any experimentation on these MEFs is 4N. Upon Cre expression, recombination between the duplicated sister chromatids yields either the same parental configuration or an inverted recombination to yield a pair of acentric and dicentric fragments (Figure 1B). The acentric fragments are not maintained during repeated cell divisions and the dicentric fragments undergo breakage-fusion-bridge cycles in subsequent mitoses and are eventually lost (Zhu et al., 2010). Since the iLoxP sites bear an externally flanking surface marker (human Cluster of Differentiation 2-hCD2), all inverted recombination events were assessed by the loss of this marker and cells were sorted by flow cytometry to obtain hCD2 Plus (parental controls) and hCD2 Minus (targeted chromosome loss) populations (Figures 1B and 1C). For brevity, we refer to cells with engineered chromosome loss events as ICLs (cells with <u>I</u>nduced-<u>C</u>hromosome <u>L</u>osses) and their unrearranged parental counterparts as controls. Previous adaptation of this strategy, in the context of studying loss of heterozygosity (LOH), has shown that recombination-mediated marker loss also correlated strongly with chromosome loss in the lymphoid system in mice (Zhu et al., 2010), suggesting that marker loss is a faithful measure of the predicted chromosome loss event both *in vitro* and *in vivo.*

**Figure1:**
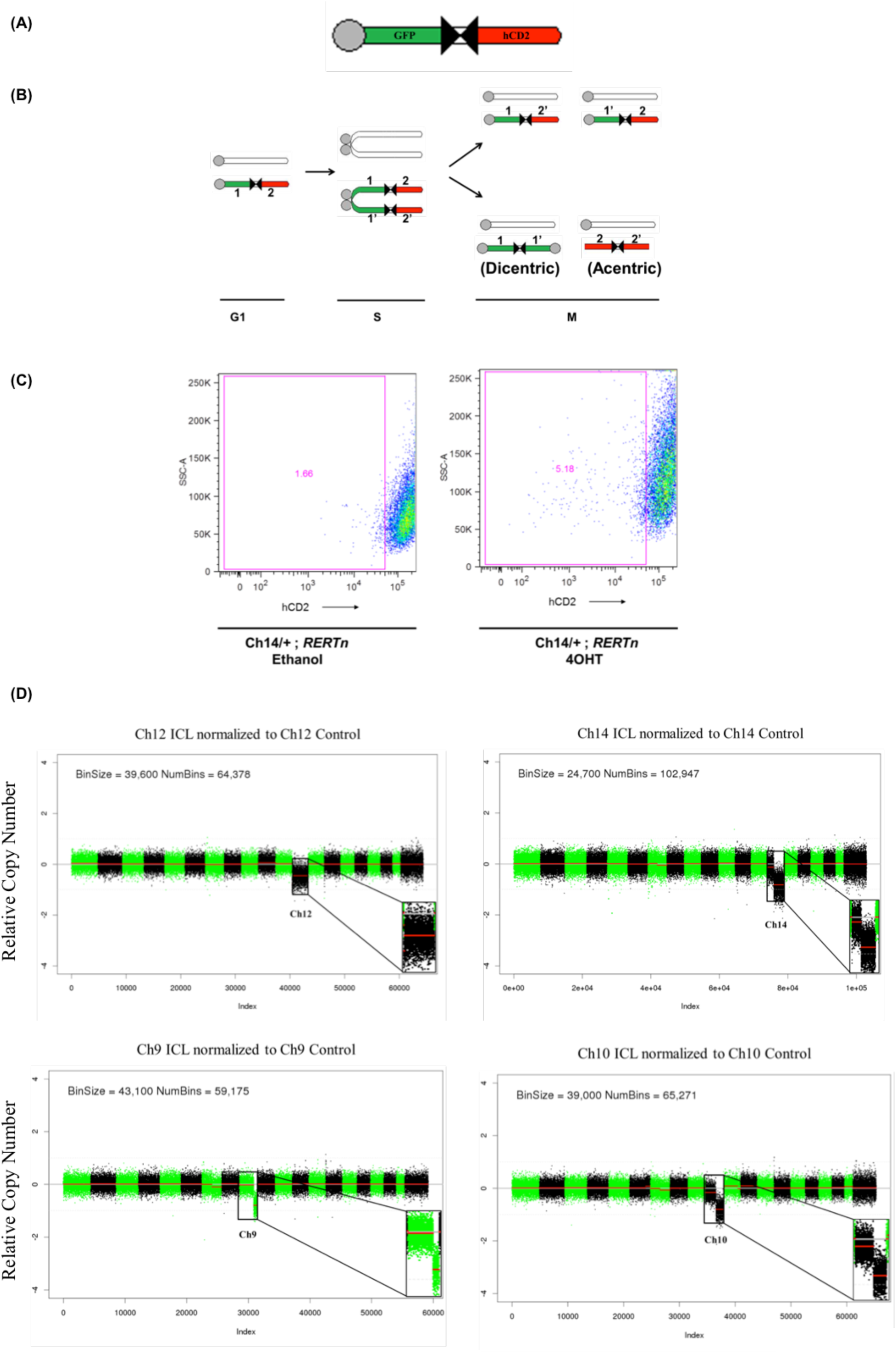
Generation of ICL in MEFs by Cre recombination. (A) Schematic of the inverted lox P (iLoxP) site, flanked by sortable GFP and hCD2 markers. (B) Model of reciprocal recombination yielding parental configurations 1/2’ and 1’/2 and dicentric/acentric configurations as a result of inverted recombination yielding configurations 1/1’ and 2/2’. (C) Representative FACS plot showing loss of hCD2 marker expression of Ch14 ICL MEFs after treatment with 4-hydroxy tamoxifen (4OHT) for 14 days to induce Cre, compared to Ethanol treated control cells. (D) Shallow whole genome sequencing (copy number profiles) of large T antigen immortalized, MEFs after exposure to Cre recombinase and sorted for control (hCD2 plus) and ICL (hCD2 Minus) cells for chromosomes 12, 14 and 9 and 10, without any *in vitro* culturing, post FACS sort.

### Validation and tumorigenic potential of early passage aneuploid cells generated by Cre-recombination

In order to demonstrate that the expected chromosome loss event took place after Cre-mediated recombination and prior to any/extensive *in vitro* expansion of the sorted cells, we performed low pass whole genome sequencing (WGS) directly on the FACS sorted ICL and control cells (without any *in vitro* culturing after the sort). As shown in Figure 1D, in each of the 4 chromosome lines sequenced, compared to the respective control lines, the ICL lines harbored a notable copy number reduction only in the in the corresponding targeted chromosomes. The absolute Z score, a measure of the number of standard deviations from the mean, was greater than 1, only for the targeted chromosomes in all the ICL lines. The Ch14, Ch9 and Ch10 ICL lines harbored a strong copy number reduction in the distal portions of the corresponding targeted chromosomes. This can be explained by the fact that the hCD2 marker (used to FACS sort), which is present on the distal portion on the chromosome (Figure 1A), is lost initially, followed by the remaining portion of the chromosome in subsequent cell cycles. Accordingly, when the ICL (and the control cells) are expanded further in culture for subsequent experiments (see next section), we observe the loss of entire chromosomes in all four lines.

We also obtained similar results on performing whole genome sequencing on the sorted ICL and control cells after minimal *in vitro* passaging. Only the targeted chromosome exhibited notable copy number reduction (Figure S1A) and the absolute Z score was greater than 1, only for the targeted chromosomes in all the ICL lines. Although, in this experiment we observed only a small copy number change in chromosome 10 in the Ch10 ICL which can be attributed to the stochastic nature and frequency of the recombination events in the different chromosomal lines at early time points. At later time points as expected, a higher proportion of the ICL cells exhibit chromosome 10 losses (see next section). Since the populations analyzed were sorted solely for the presence or absence of the hCD2 marker, we anticipated at very low passage numbers, very few differences outside the ICL event would be observed. Indeed, as shown in Figure S1A, compared to the control cells, the targeted chromosome loss was the only prominent event in the ICL cells.

All 4 of these low passage ICL lines were injected into immuno-compromised, athymic nude mice to determine the tumorigenic potential compared to the control populations. In 2 of the 4 chromosome lines tested (Ch9 and Ch14), the ICL cells showed dramatic enhancement of tumorigenic potential (Figure 2A). Importantly, neither of these ICL lines showed enhanced growth potential *in vitro*, suggesting that the advantages conferred on the ICL cells by the targeted chromosome loss are manifested only in certain contexts, such as the stressed environment in the mice (Figure S1B). Thus, our Cre-recombination mediated strategy was robust in generating the loss of the targeted chromosomes in MEFs and in 2 out of 4 early passage ICL lines, the targeted chromosome loss is largely sufficient to promote tumor formation in nude mice.

**Figure 2:**
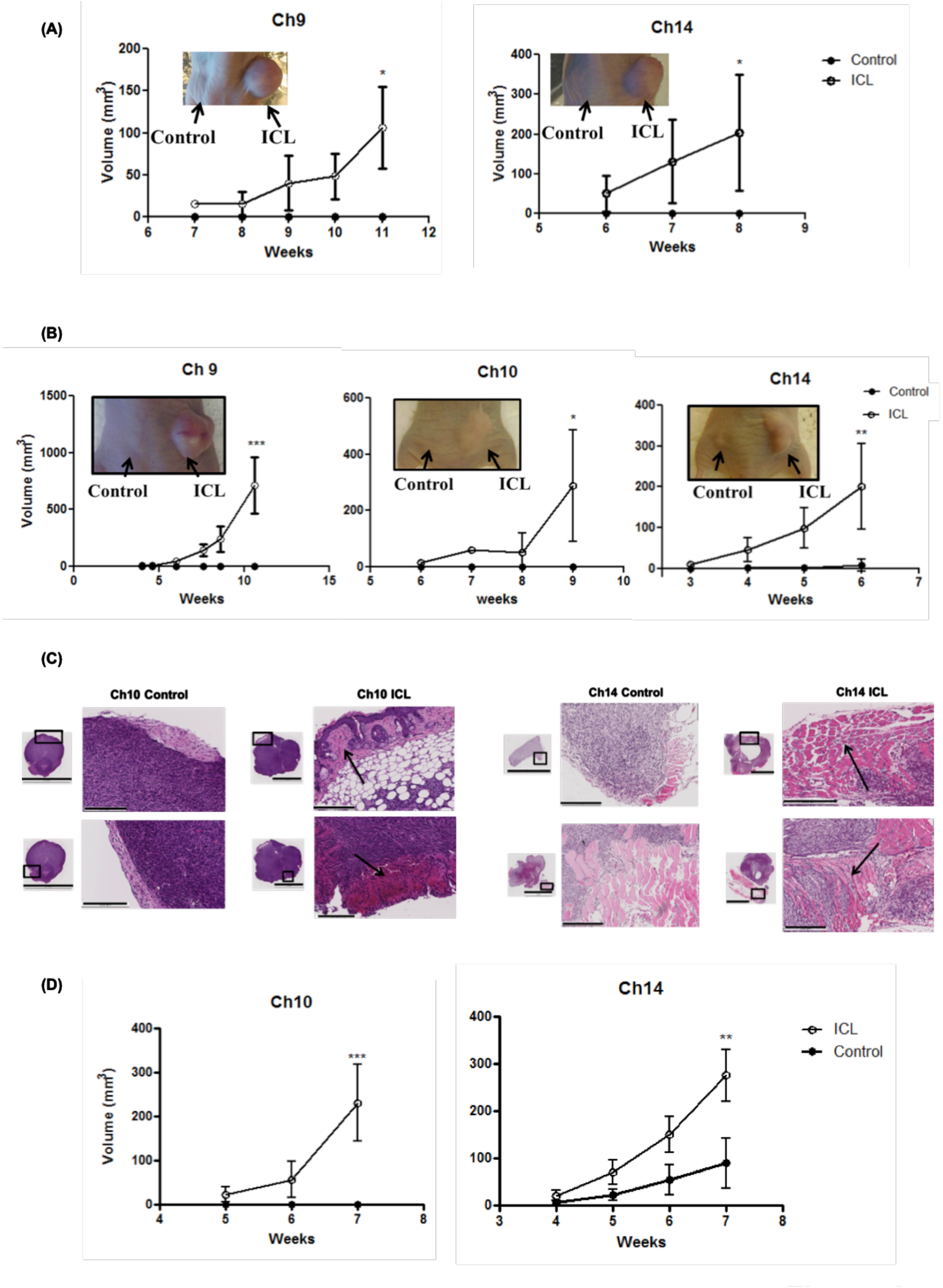
Enhanced tumor formation in immunocompromised mice. (A) Tumor growth curve after early passage ICL cells were injected into flanks of athymic nude mice (n = 5 per group and error bars denote SEM, p<0.05 for Ch9 and Ch14). (B) Tumor growth curve after later passage ICL cells were injected into flanks of athymic nude mice (n = 5 per group and error bars denote SD, p<0.0005 for Ch9, p<0.05 for Ch10 and p<0.005 for Ch14). (C) Tumors sections were stained with H&E to assess tumor histology (scale bars denote 5mm for the inset and 200μm for the zoomed image) Arrows in the ICL tumors sections indicate regions where tumors have invaded into the adjacent tissue. (D) Tumor growth curve after later passage ICL cells were injected into flanks of NOD/SCID mice (n = 5 per group and error bars denote SD p<0.0005 for Ch10 and p<0.005 for Ch14).

### Characterization, tumorigenic potential and *in vitro* growth properties of later passage ICL cells

Prior to all subsequent experiments, the immortalized MEFs were sorted three times, serially, to improve the purity of hCD2 Plus (control) and Minus (ICL) populations. As anticipated for tetraploid cells lacking pocket protein activity, whole chromosome gains and losses were increased during serial passaging both in the ICL and control cell lines. Importantly, karyotyping analysis of these “later passage” MEFs indicated that all ICL lines showed significant levels of target chromosome loss in 70-80% of the cells (Figures S2A and S2B). Many of the non-targeted chromosomal aberrations (deviations from a copy number of 4) observed in the parental control cells are also found in the respective ICL lines, indicating that the control and ICL lines suffer some common changes that may result from cell passage pressures (see arrows and asterisks in Figure S2A). Loss of the target chromosome was by far the most prevalent change in the ICL cells (~80% of the metaphases) (Figure S2B).

In order to determine if passaging influenced the tumorigenic potential of these later passage lines we repeated the xenograft transplantation experiments into immunocompromised mice. ICL lines for Ch9 and Ch14 retained their tumorigenic potential and ICL for Ch10 now gained tumorigenic potential as well (Figure 2B). ICL lines (Ch9, 10 and 14) formed large tumors, which exhibited pronounced neural, muscular and vascular invasiveness, whereas controls either did not grow at all (Ch9) or formed small, non-invasive lesions (Ch10 and 14) (Figure 2B, C). This result was corroborated for the Ch10 and Ch14 lines, in the more fully immunodeficient NOD/SCID mouse model, to rule out any immune component to the failure of control cells to form tumors (Figure 2D). Tumors derived from ICL lines still had <4 copies of the targeted chromosome in ~85% of the cells karyotyped (example shown in Figure S3A).

There are multiple reasons that we attribute the tumor phenotype we observe to the loss of the targeted chromosome and not to other sporadic basal chromosome variations in these cells. First, in whole genome sequencing of our early passage MEFs, the only prominent, initial aneuploidy event that partitioned with the tumor promoting phenotype was the reduction in the copy number of the targeted chromosome in the ICL lines (Figure S1A). Second, since each ICL line and matched control was derived from a common MEF line, most of the non-targeted chromosome variations were present in both the ICL and the controls (Figure S2A and S2B). Also, unlike the non-targeted chromosomal variations, the targeted chromosome losses in the ICL lines were the most penetrant events occurring in about 70 to 80% of the ICL cells. Third, there was no clonal selection for any other predominant chromosomal abnormality during the course of tumor formation (data not shown). Finally, similar results were obtained when the experiments were performed using 2 to 4 different MEF lines (biological replicates) for any given ICL chromosome, and we observed the pro-tumorigenic phenotype for three different ICL chromosome lines (9, 10 and 14) (Figure 2B). Collectively, these data indicate that, later passage ICLs of mouse chromosomes 9, 10 and 14 results in a significant increase in the *in vivo* tumorigenic potential.

As seen with the early passage cells, the later passage growth rate in cell culture was either the same or lower in the ICLs versus controls. Three out of our four ICL lines (Ch10, 12, 14) grew significantly slower *in vitro* relative to their controls, suggesting reduced cellular fitness (Figure 3A and Figure S3B). The fourth ICL line missing Ch9 had no significant change in growth rate (Figure S3B). ICL cells also formed fewer colonies when plated at very low seeding, further implying an *in vitro* growth defect (Figure 3B). Thus, expansion of the ICL lines in culture maintained or enhanced the tumorigenic potential of the lines which had initially lost a single chromosome and similarly did not affect the relative growth rates in cell culture. All ICL lines showed a significant growth advantage over the controls under anchorage-independent conditions (Figure 3C), a property consistent with their enhanced tumorigenic potential. When the ICL lines were explanted from the tumors, they did not retain the proliferative advantage (seen *in vivo*) highlighting that the growth advantage is manifested only under conditions requiring anchorage independence (Figure S3C).

**Figure 3:**
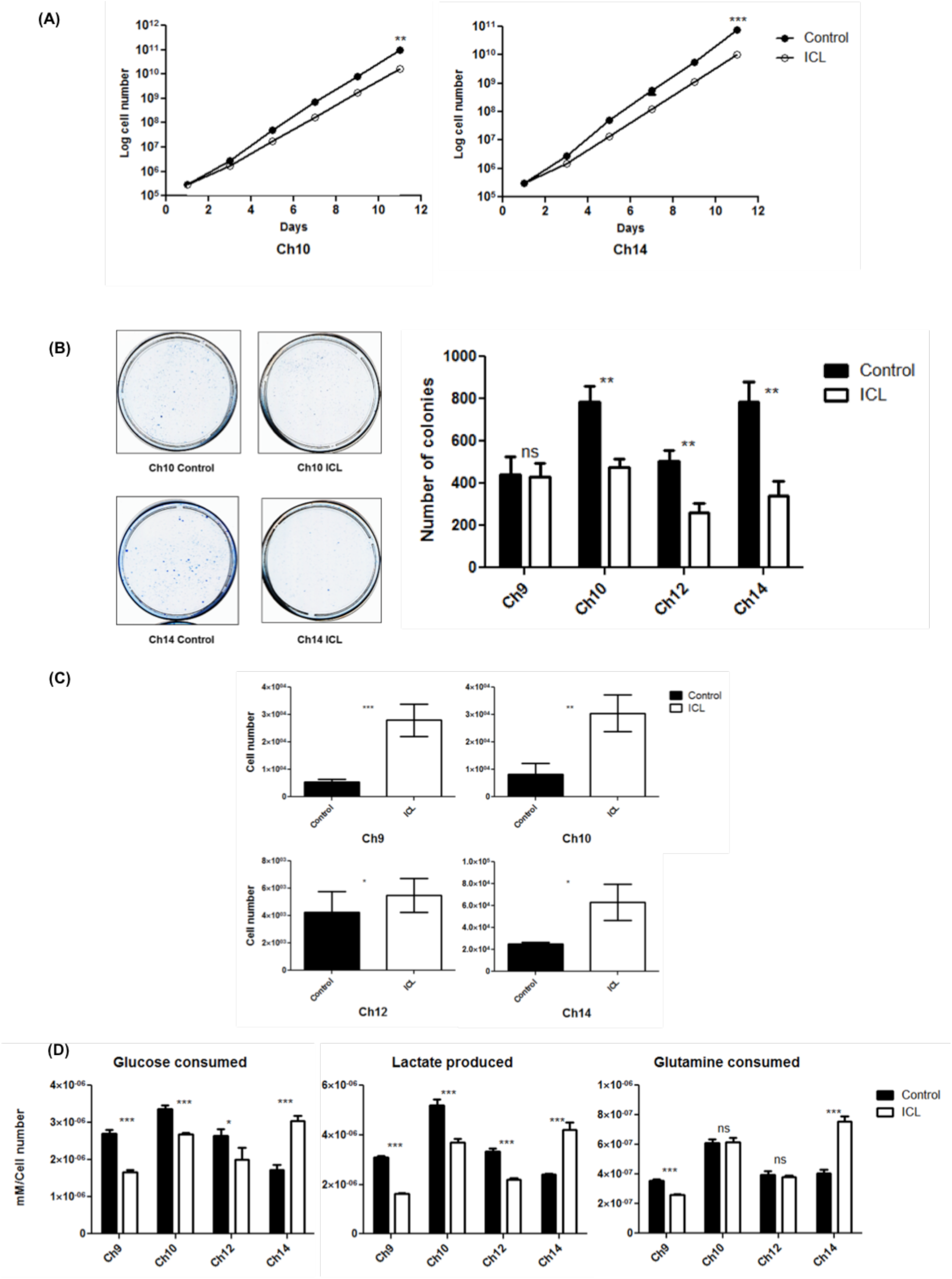
*In vitro* characterization of later passage ICL MEFs. (A) Growth curve of the later passage ICL lines Ch10 and Ch14 under adherent *in vitro* culture conditions (n=3 for each data set and error bars denote SD, p<0.005 for Ch10 and p<0.0001 for Ch14). (See also Figure S3B) (B) Colony formation in ICL and control MEFs. 5000 cells were seeded in 10 cm plates and stained with Methylene blue after three days. Representative colony formation images in Ch10 and Ch14 are shown (left) and quantification of colonies formed (right) in the 4 ICL lines (n=3 for each data set and error bars denote SD, p<0.005 for Ch10, Ch12 and Ch14 and ns for Ch9). (C) Quantification of growth in ultra-low adherent, sphere forming conditions for the 4 ICL lines (n=3 for Ch10 and Ch14, n=6 for Ch9 and Ch12 and error bars denote SD, p<0.0001 for Ch9, p<0.001 for Ch10 and p<0.05 for Ch12 and Ch14). (D) Analysis of metabolites (glucose consumed, lactate produced and glutamine consumed) for the ICL lines. Spent media was analyzed after each ICL & control line was grown in culture for three days. Media in identical culture conditions, but with no plated cells, was used as a baseline for all samples.

In order to determine if metabolic changes characteristic of certain tumors and trisomic MEFs were also present in our ICL cells, we performed metabolite analysis on the conditioned media from these cells (Williams et al., 2008). Ch14 ICL cells had increased levels of lactate production and glucose consumption in comparison to the Ch14 controls (Figure 3D). These altered metabolic phenotypes were not manifested in Ch9, Ch10 and Ch12 ICL cells, suggesting that metabolic changes are likely chromosome-specific effects.

Taken together, given that the tumor phenotypes of at least two ICL lines (9 and 14) were observed prior to the accrual of any non-targeted chromosomal anomalies, argues strongly that these targeted chromosome losses were the driving event in the acquisition of tumorigenicity. We cannot rule out the possibility that Ch10 ICL, which acquired tumor potential only after longer passaging, had suffered other events that facilitated their transformation. We suspect that the initial loss of Chr10 must at least be contributing to the tumor phenotype since, in the 4 different Ch10 lines examined, only the ICLs (and never the control cell lines), after extensive passaging, ever acquire tumorigenic potential.

### Effects of tetraploidy and chromosome identity on tumorigenecity

ICL of Ch12 consistently did not produce tumors, arguing against the possibility that any breakage and rejoining events per se that take place during the process of chromosome loss transforms immortalized MEFs. Also, the basal aneuploidy arising from the tetraploidization and/or p53/pRb attenuation, observed in the parental control lines was not sufficient to form tumors in the mice, suggesting a critical role for the targeted chromosome loss in tumorigenesis. We note that tetraploidization of our ICL lines may be required to manifest the tumorigenic potential upon chromosome loss. p19 immortalized, diploid, MEFs with targeted chromosome losses (Ch10 and 14) did not produce tumors (data not shown). In addition, to test the effects of chromosome loss in an independent diploid model *in vivo*, a knock-in allele of Id1 driving tamoxifen inducible CRE recombinase was employed in the Ch10 and Ch14 iLoxP mice (Nam and Benezra, 2009). Id1 is expressed in long-term repopulating hematopoietic stem cells (Jankovic et al., 2007) and other stem cell populations (Nam and Benezra, 2009). While ~50% of the blood cells in these mice exhibited hCD2 marker loss at 30 days post Cre induction, at a later time point (80 days), this number dropped to 17%, indicating that the monoploid cells have a proliferative disadvantage in comparison to the diploid population (Figure S4A). This suggests that the effects of inducing chromosome loss can have different effects depending on the original ploidy and the tissue type where aneuploidy is induced. It is noteworthy that in the vast majority of human tumors with numerical chromosome losses, the modal chromosome number is >2N (both on an individual chromosome basis, and for any chromosome in general) (Figures S4B and S4C), implying that the tetraploid state is a clinically relevant background to assess the contribution of chromosome losses to tumorigenesis

The variation in tumorigenic properties and tumor latency between the ICL lines suggests that there may be chromosome-specific effects with regards to tumorigenesis. Highlighting this, Davoli *et al* used a computational method to assign ‘CHROM’ scores to human chromosomes based on the mutational signatures of 300 highest ranked tumor suppressor genes (TSGs) and 250 oncogenes (OGs) and observed that in human tumors, chromosomes with a higher CHROM score (high density of top ranked TSGs) were lost more frequently (Davoli et al., 2013). We mapped these human TSGs and OGs onto mouse chromosomes and observed that while tumor suppressor genes from these ranked lists were present on Ch14 (high density of top ranked TSGs) and Ch9 (moderate density), Ch10 and 12 were largely devoid of highly ranked TSGs (Table S1). While higher density of potent TSGs for Ch9 and Ch14 would be expected to be associated with increased tumorigenic potential upon chromosome loss, a lack of highly ranked TSGs by itself did not result in failure to form tumors, as demonstrated in the Ch10 ICL cells, which readily formed tumors. Thus, both general chromosome loss effects, as well as chromosome-specific effects, likely contribute to tumorigenic potential in this model system.

### ICL induced genomic instability and DNA damage

In an attempt to understand the mechanism by which the ICL lines became inherently better suited for anchorage independent growth (and by inference tumor growth), we analyzed their ability to undergo an epithelial-to-mesenchymal transition (EMT) more rapidly than parental controls or activate pro-proliferative pathways and/or autophagy, as these are common mechanisms by which cells attain anchorage independence (Guadamillas et al., 2011). We found no clear changes in these measures suggesting that other mechanisms are driving the observed phenotypes (data not shown). Aneuploidy has been shown to potentially cause a mutator phenotype that further induces more genomic instability, which could facilitate adaptation to the stresses of anchorage independent growth. In haploid yeast strains, for example, addition of an extra chromosome caused genomic instability phenotypes like chromosome mis-segregation errors and increased mutation rate (Sheltzer et al., 2011). In *S. cerevisiae*, aneuploidy caused by Hsp90 inhibition enabled yeast strains to adapt well to other unrelated stress conditions by inducing karyotypic diversity (Chen et al., 2012). Additionally, trisomic MEFs with certain oncogenic activations, acquired new karyotypic changes at later passages that provided better adaptive fitness to these cells (Sheltzer et al., 2017).

For these reasons we sought to determine if the induced aneuploid cells had evidence of further genomic instability beyond target chromosome loss. Karyotyping analysis revealed increased chromosome instability in all later passage ICL lines *in vitro*, demonstrated by increased average chromosome loss per metaphase relative to the control counterparts (6.2 vs 4.2 for Ch9, p<0.005; 12.3 vs 5.7 for Ch10, p<0.0001; 4.8 vs 2.9 for Ch12, p<0.05 and 11.8 vs 6.6 for Ch14, p<0.0001), average structural rearrangements per metaphase (5.3 vs 1.5 for Ch9, p<0.0001; 14.2 vs 8.5 for Ch10, p<0.0001; 2 vs 1.1 for Ch12, p<0.05 and 11 vs 6.7 for Ch14, p<0.05) and percentage of metaphases with marker chromosomes (chromosome fragments) (90 vs 75 for Ch9,ns; 80 vs 20 for Ch10, p<0.0005 and 35 vs 15 for Ch12, ns) (Figure 4A). Also, compared to the controls, we observed an increase in basal γH2AX foci per nuclei (a read-out of DNA double strand breaks) in ICL cells (124 vs 92 in Ch9, p<0.0001; 102 vs 53 in Ch10, p<0.0001; 97 vs 67 in Ch12, p<0.0001; 78 vs 63 in Ch14, p<0.05) (Figure 4B). The increase in γH2AX foci in the ICL cells was also observed when the cells were grown in the low adherence sphere forming conditions (73 vs 47 in Ch10; p<0.0001 and 75 vs 47 in Ch14, p<0.0001) (Figure S5A). Surprisingly, we noticed that the ICL lines exhibited a modest yet significant increase in the percentage of cleaved caspase positive cells, an apoptosis marker, despite not displaying signs of cell death (13.1 vs 0.8% in Ch9, p<0.0001; 2.0 vs 1.0 % Ch10, p<0.05; 1.3 vs 0.9% in Ch12, p<0.05 and 0.7 vs 0.2% in Ch14, p<0.0005) (Figure S5B). This may also contribute to the enhanced mutator phenotype. Similarly, tumors derived from ICL lines showed increased numbers of γH2AX positive cells in comparison to the controls (4.4 vs 1.8 % in Ch10, p<0.0001 and 3.7 vs 2.4% in Ch14, p<0.05) (Figure 4C). Together, these data indicate that the later passage ICL lines are associated with high levels of chromosomal instability and DNA damage and this mutator phenotype could potentially allow the ICL cells to sample more karyotypic space. We recognize that this mutator phenotype is not sufficient for transformation, as Ch12 ICL cells did not form tumors despite having elevated DNA damage. Thus, while the mutator phenotype may be required to enable the ICL cells to adapt well to stress conditions (such as enhanced growth in substratum-free conditions and in the subcutaneous space of mice), there may be other mechanisms intrinsic to chromosome loss that are required for tumorigenesis.

**Figure 4:**
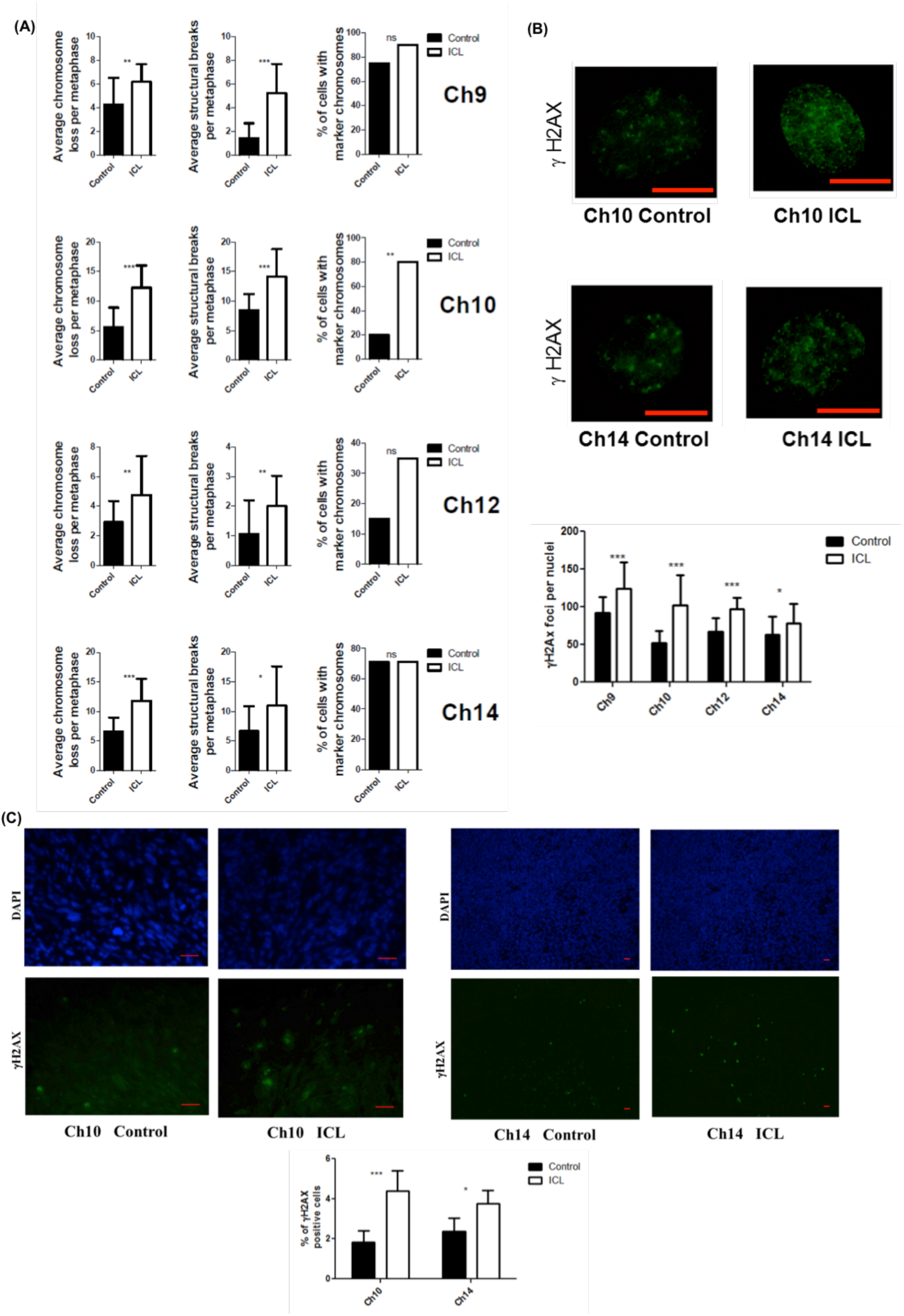
Increased chromosomal instability and DNA damage in ICL *in vitro* lines and tumors. (A) Chromosomal instability readouts, of later passage lines, assessed by average chromosomes loss per metaphase, average number of structural rearrangements per metaphase and percentage of metaphases with marker chromosomes (n=20 for each dataset and error bars denote SD). (B) Determination of number of γH2AX foci in ICL and control nuclei and quantification (scale bars denote 10μm, p<0.0001 for Ch9, Ch10 and Ch12, p<0.05 for Ch14; n=25 for each dataset and error bars denote SD). (See also Figures S5A and S5B) (C) Number of γH2AX positive cells in formalin fixed ICL and control tumor sections by immunofluorescence staining and quantification (Scale bars denote 20μm, p<0.0001 for Ch10 and p<0.05 for Ch14).

## Discussion

In this study, we have successfully adapted the iLoxP inverted recombination system to generate an aneuploidy model, inducing targeted chromosome losses in the context of a tetraploid state, without perturbing the expression of a specific protein. Variations of this method have also been used previously to introduce various genomic aberrations in mice (Gregoire and Kmita, 2008; Lewandoski and Martin, 1997; Zhu et al., 2010). We found that ICL of mouse chromosomes, while not providing a substantial growth advantage under normal *in vitro* conditions, facilitated growth under the stressful conditions of anchorage-independent growth and *in vivo* tumor formation. The parental controls in later passages, which exhibited a minor degree of aneuploidy independent of the targeted chromosome loss, showed no or reduced growth in immunocompromised mice compared to matched ICL lines. This observation was consistent across multiple separate ICL lines for a given chromosome, as well as across multiple chromosomes. Further, in early passage MEFs, the only pronounced copy number change was in the targeted chromosomes, and in 2 out of the 4 chromosome lines tested, this targeted change alone was sufficient to drive tumorigenesis. Taken together, these results demonstrate that the initial loss of the targeted chromosome is critical for these tumor phenotypes. We note that recent data suggests that aneuploid cells may thrive in human tumors due to escape from immune surveillance (Davoli et al., 2017). Our data in immuno-compromised mouse models argues for an additional immune-independent growth advantage of aneuploid cells.

While there are deviations from the baseline copy number of 4 in a few chromosomes in the later passage control lines, we speculate that the reasons that these aberrations are not sufficient to initiate *in vivo* tumors are: 1) these baseline changes in the control lines are not as penetrant as the targeted losses in the ICL lines in the population and therefore the ensuing instability is below the threshold to similarly accelerate tumorigenesis and/or 2) the reduction to copy number of 2 is necessary to manifest the tumor phenotype (which is observed only in the ICL cells). We also observed that the initial whole-chromosome loss, irrespective of the identity of the lost chromosome, induced a significant increase in chromosomal instability, DNA damage and secondary chromosome losses. This feature could enable the ICL cells to sample more genomic permutations and enable them to thrive under stressful, substratum free conditions. Thus, in addition to maintaining the loss of the targeted chromosome, the ICL cells accumulated other chromosomal changes more rapidly than the controls. Other models of whole chromosome aneuploidy also generate a mutator phenotype, causing further chromosomal instability and DNA damage (Crasta et al., 2012; Janssen et al., 2011a; Sheltzer et al., 2011). In addition, DNA damage has been shown to increase chromosome instability through the stabilization of kinetochore-microtubule interactions (Bakhoum et al., 2014) thereby forming a possible feed forward loop of chromosomal instability. Whether enhanced chromosome mis-segregation observed in the ICL lines is a cause or consequence of enhanced DNA damage and how this cycle is started by the initiating chromosome loss event will require further study.

After the inverted Cre recombination, the dicentric fragments, which undergo BFB cycles, could invade and integrate randomly into other regions of the genome (instead of being lost) thereby promoting genomic instability and tumorigenesis. BFB cycles of dicentric fragments are a common cause of chromosome loss in cancer cells and such events have been shown to induce genomic redistribution, genomic instability and tumor heterogeneity in a variety of tumor types (Artandi et al., 2000; Gisselsson et al., 2000; Selvarajah et al., 2006). Dicentric fragments can also result in chromothripsis, a catastrophic chromosome shattering and reorganization phenomenon. In this model, instead of being broken, the dicentrics result in the formation of chromatin bridges, leading to nuclear envelope rupture and DNA repair, and eventually to structurally reorganized chromosomes (Maciejowski et al., 2015). Mis-segregating chromosomes (that lead to whole chromosome losses in the daughter cells) have also been shown to trigger gross genomic redistributions. The mis-segregating chromosomes induce micronuclei formation, where they are pulverized and then integrated back into the genome (Crasta et al., 2012). Mis-segregating chromosomes also produce structural chromosomal changes by being trapped and damaged in the cleavage furrow during cytokinesis, leading to DNA breaks and unbalanced translocations (Janssen et al., 2011b). Thus two processes, pure chromosome loss events and structural genomic rearrangements, are intimately connected. Though we cannot distinguish between the contributions of these two processes to transformation, our model to induce chromosome loss via the formation of dicentrics nonetheless attempts to faithfully mimic the actual evolution of cancer cells.

We note that the tumorigenic potential of the ICL lines are manifested only in a tetraploid background. This benefit of chromosome losses from a tetraploid genome is consistent with previous data from other groups. Tetraploid mouse cells have been shown to lose chromosomes when they form tumors in mice, suggesting that tetraploidy may be a state where chromosome loss is better tolerated or even beneficial (Davoli and de Lange, 2012). p53^-/-^ tetraploid mouse mammary epithelial cells were more prone to whole chromosome aneuploidies than their diploid counterparts and the tetraploidization promoted tumor formation of these cells in nude mice (Fujiwara et al., 2005). Also, it has been shown, using *in silico* analyses, that a near triploid state endows the maximum fitness to the adaptation of the cancer cells (Laughney et al., 2015). Importantly, in this study, the tetraploid state (rather than the diploid state) has a shorter barrier to reach the near triploid state. In addition, Carter et al, using a computational method, showed that whole genome doublings are common occurrences in human tumors and that the evolved karyotype of these tumors is a result of losing chromosomes after the initial doubling (as opposed to sequential gaining of chromosomes on a diploid background) (Carter et al., 2012). Thus, these studies highlight the importance of passing through a tetraploid state followed by chromosome losses in tumorigenesis.

In summary, our ICL model demonstrates that individual chromosome loss events in a tetraploid background can be a potent driver of tumorigenesis in mouse cells and provides a new platform to further our understanding of the consequences of whole chromosome loss events in cancer.

## Materials and Methods

### Mouse embryonic fibroblasts (MEFs) FACS sorting

Primary mouse embryonic fibroblasts from Ch9, 10, 12 and 14 mouse lines were isolated from embryonic day 13.5 old mice and were immortalized by transfecting with SV40 Large T antigen. Recombination between the iLoxP sites was induced in Ch10 and Ch14 immortalized MEFs, which expressed a ubiquitous tamoxifen-responsive Cre-ER (RNA PolII CreER, *RERT*), by treating with 1μM 4-hydroxy tamoxifen (4OHT) for 14 days. Recombination was induced in Ch9 and Ch12 immortalized MEFs by transducing with an adenovirus expressing Cre recombinase for 18h. Recombined MEFs were stained with an hCD2 antibody (BD Pharmingen) and flow sorted to obtain hCD2 plus (control) and hCD2 minus (ICL) populations.

### Growth rates, Seeding efficiency and Metabolite analysis

For determining the growth rates, 300,000 cells were plated in triplicate in 60 mm tissue culture dishes (or 50,000 cells in each well of 6 well plates). After every two days, cells were trypsinized, counted and re-plated at the starting density. For seeding efficiency measurements, 5000 cells were plated in 100mm culture dishes. After three days, cells were stained with 0.05% Methylene blue and the colonies were counted. For metabolite analysis, cells were plated as described for growth curves. After three days, growth media was collected, spun down to remove debris and residual cells and analyzed using an YSI 7100 MBS Mutliparameter Bioanalytical System to obtain metabolite concentrations for glucose, lactate and glutamine. Cell culture media from plates exposed to identical culture conditions, but without plated cells was used as the baseline. Two tailed unpaired t test was used to determine statistical significance.

### Growth in immunocompromised mice

ICL cells and their respective controls were injected into the flanks of athymic nude mice at a cell density of 250,000 in 200 μl phosphate buffer for later passage Ch10, Ch14 cells; 1,000,000 in 200 μCh12 cells; 5,000,000 in 200l phosphate buffer for later passage Ch9, Ch12 cells; 5,000,000 in 200 μl phosphate buffer for early passage Ch14 cells and 15,000,000 in 200 μl phosphate buffer for early passage Ch9 cells. These mice were monitored for tumor formation and tumor dimensions were measured weekly. The tumor volume was calculated using the formula Volume=0.5∗Length∗(width)^^^2. Two tailed unpaired t test was used to determine statistical significance. All the immunocompromised mice were ordered from Taconic Biosciences. Housing regulations and experimentation on the mice were approved by the MSKCC Institutional Animal Care and Use Committee (IACUC) and Research Animal Resource Center (RARC).

### Karyotypes and Shallow whole genome sequencing

For metaphase spreads, one million cells were plated in a 10 cm dish, and after 24 hours treated with 0.2μM colcemid for 4 hrs. Cells were trypsinzed, incubated in 75mM KCl and fixed with 75% methanol: 25% acetic acid mixture for 5 mins. After two additional washes with fixative, cells were dropped from a height of about 2 inches on pre-chilled glass slides, which were held over a water bath. Slides were allowed to dry overnight and stained with 0.02% Giemsa and submitted for karyotype analysis. Images were analyzed using Applied Spectral Imaging. Fisher exact test was performed to calculate statistical significance of chromosome number variation between the controls and the ICL metaphases and a false discovery correction was applied to the p values. For the chromosomal instability readouts, two tailed unpaired t test was used to determine statistical significance for average chromosome loss and structural breaks per metaphase. A two tailed Z test was used to determine statistical significance for the percentage of cells with marker chromosomes. Shallow whole genome sequencing was carried out at the Integrated Genome Operations core facility at MSKCC. 50ng of purified DNA from each control and ICL line was sequenced on Illumina HiSeq2000 (10 million reads per sample with 100mer paired end sequencing). The sequencing results were aligned to mouse mm9 genome and copy number changes were estimated in 300kb bins.

### Regrowth in culture and karyotyping of tumors

Tumor-bearing mice were euthanized and the tumors were excised from the flanks, dissociated by incubation with papain (500 units), vigorously triturated and incubated for 15min at 37°C. The cell suspension was centrifuged and resuspended in an ovomucoid protease inhibitor media solution to inhibit residual papain, followed by vigorous trituration. Cells were spun again and the pellet was resuspended in cell culture media and plated in 6-well plates. Growth and karyotyping analysis were performed as described above.

### Growth in Ultra Low Adherence Conditions

For this assay, 20,000 cells were plated on Ultra low attachment 24-well plates (Corning Costar). After three days, cells were pelleted at 1000rpm for 10min and washed with STE buffer (100mM NaCl, 10mM Tris pH7.4, 1mM EDTA). Cells were stained with 0.4% trypan blue solution, and live cells were counted.

### Immunofluorescence

ICL and control cells were plated in chamber slides, fixed with either methanol/acetone (for γH2AX) or 4% paraformaldehyde (for cleaved caspase) and stained with the primary antibodies(γH2AX – JBW301, EMD Millipore; Cleaved Caspase - #9661, Cell signaling). For measuring DNA damage in the spheres formed in low adherent conditions, the spheres were harvested as described above and treated with hypotonic 75mM KCl solution at 37°C for 15 minutes and then cytospun onto glass slides and immunofluorescence was carried out as above.

### Blood analysis

6 week old mice harboring the iLoxP construct and a tamoxifen inducible Cre recombinase expressed in long term repopulating hematopoietic stem cells (*Id1*^*IRES*-*creERT2*^) were injected with 16μg of Tamoxifen in corn oil over 4 days. 30 days and 80 days post tamoxifen administration, 100μL of tail blood was collected from these animals and stained with the hCD2 antibody (described earlier). The red blood cells were next lysed with BD FACS Lyse solution and the remaining lymphocytes were analyzed using flow cytometry for the proportion of cells that had lost the hCD2 marker.

### CHROM analysis

CHROM analysis on mouse chromosomes was done based on the methods described previously for human chromosomes by Davoli et al (Davoli et al., 2013). Briefly, the chromosomal loci for the top 300 human tumor suppressor (TSGs) and 250 oncogenes (OGs) used in their analysis were mapped to the mouse genome using the UCSC genome browser while maintaining the rankings for each gene.

### Analysis of the human tumors from the Mitelman database

Human solid tumors that had lost (or gained) any chromosome were retrieved from the Mitelman database (http://cgap.nci.nih.gov/Chromosomes/Mitelman) and sorted according to the modal chromosome counts. This set was then stratified into either the group whose modal chromosome count was >51 (>2N) or the group whose modal chromosome count was between 36 and 50 (~2N). The total number of cases in each of these groups was then quantified and the proportion of cases within the group (>2N or ~2N) that had lost (or gained) was calculated. For the analysis involving tumors that have lost (or gained) specific chromosomes, a similar approach was used with the exception of retrieving the tumors that have lost (or gained) the corresponding chromosome. Two tailed 2 sample t test to compare sample means, was used to calculate statistical significance.

## Acknowledgments

The authors would like to thank Y.Zhuang for generously providing the Ch10 and Ch14 iLoxP mouse lines, Juan-Manuel Schvartzman and Dr. David Pellman (Harvard Medical School) for critical reading of the manuscript, MSKCC Flow Cytometry core facility for FACS assistance, Molecular Cytogenetics core facility for the karyotyping analysis, the Integrated Genomics Operation (IGO) core facility for the whole genome sequencing and analysis, and J.Cross from the Cancer metabolism center core facility for performing the metabolite analysis. This work was supported by a grant from the Geoffrey Beene Cancer Research Center and NIH/NCI Core Grant P30 CA008748

## References

Artandi, S.E., Chang, S., Lee, S.L., Alson, S., Gottlieb, G.J., Chin, L., and DePinho, R.A. (2000). Telomere dysfunction promotes non-reciprocal translocations and epithelial cancers in mice. Nature 406, 641-645.

Bakhoum, S.F., Kabeche, L., Murnane, J.P., Zaki, B.I., and Compton, D.A. (2014). DNA-damage response during mitosis induces whole-chromosome missegregation. Cancer Discov 4, 1281-1289.

Carter, S.L., Cibulskis, K., Helman, E., McKenna, A., Shen, H., Zack, T., Laird, P.W., Onofrio, R.C., Winckler, W., Weir, B.A., et al. (2012). Absolute quantification of somatic DNA alterations in human cancer. Nat Biotechnol 30, 413-421.

Chen, G.B., Bradford, W.D., Seidel, C.W., and Li, R. (2012). Hsp90 stress potentiates rapid cellular adaptation through induction of aneuploidy. Nature 482, 246-250.

Crasta, K., Ganem, N.J., Dagher, R., Lantermann, A.B., Ivanova, E.V., Pan, Y., Nezi, L., Protopopov, A., Chowdhury, D., and Pellman, D. (2012). DNA breaks and chromosome pulverization from errors in mitosis. Nature 482, 53-58.

Davoli, T., and de Lange, T. (2012). Telomere-driven tetraploidization occurs in human cells undergoing crisis and promotes transformation of mouse cells. Cancer Cell 21, 765-776.

Davoli, T., Uno, H., Wooten, E.C., and Elledge, S.J. (2017). Tumor aneuploidy correlates with markers of immune evasion and with reduced response to immunotherapy. Science 355.

Davoli, T., Xu, A.W., Mengwasser, K.E., Sack, L.M., Yoon, J.C., Park, P.J., and Elledge, S.J. (2013). Cumulative haploinsufficiency and triplosensitivity drive aneuploidy patterns and shape the cancer genome. Cell 155, 948-962.

Duijf, P.H., Schultz, N., and Benezra, R. (2013). Cancer cells preferentially lose small chromosomes. International journal of cancer Journal international du cancer 132, 2316-2326.

Fujiwara, T., Bandi, M., Nitta, M., Ivanova, E.V., Bronson, R.T., and Pellman, D. (2005). Cytokinesis failure generating tetraploids promotes tumorigenesis in p53-null cells. Nature 437, 1043-1047.

Gisselsson, D., Pettersson, L., Hoglund, M., Heidenblad, M., Gorunova, L., Wiegant, J., Mertens, F., Dal Cin, P., Mitelman, F., and Mandahl, N. (2000). Chromosomal breakage-fusion-bridge events cause genetic intratumor heterogeneity. P Natl Acad Sci USA 97, 5357-5362.

Gregoire, D., and Kmita, M. (2008). Recombination between inverted loxP sites is cytotoxic for proliferating cells and provides a simple tool for conditional cell ablation. P Natl Acad Sci USA 105, 14492-14496.

Guadamillas, M.C., Cerezo, A., and del Pozo, M.A. (2011). Overcoming anoikis - pathways to anchorage-independent growth in cancer. J Cell Sci 124, 3189-3197.

Hein, J., Boichuk, S., Wu, J., Cheng, Y., Freire, R., Jat, P.S., Roberts, T.M., and Gjoerup, O.V. (2009). Simian virus 40 large T antigen disrupts genome integrity and activates a DNA damage response via Bub1 binding. Journal of virology 83, 117-127.

Jankovic, V., Ciarrocchi, A., Boccuni, P., DeBlasio, T., Benezra, R., and Nimer, S.D. (2007). Id1 restrains myeloid commitment, maintaining the self-renewal capacity of hematopoietic stem cells. P Natl Acad Sci USA 104, 12601265.

Janssen, A., van der Burg, M., Szuhai, K., Kops, G.J., and Medema, R.H. (2011a). Chromosome segregation errors as a cause of DNA damage and structural chromosome aberrations. Science 333, 1895-1898.

Janssen, A., van der Burg, M., Szuhai, K., Kops, G.J.P.L., and Medema, R.H. (2011b). Chromosome Segregation Errors as a Cause of DNA Damage and Structural Chromosome Aberrations. Science 333, 1895-1898.

Laughney, A.M., Elizalde, S., Genovese, G., and Bakhoum, S.F. (2015). Dynamics of Tumor Heterogeneity Derived from Clonal Karyotypic Evolution. Cell Rep 12, 809-820.

Lewandoski, M., and Martin, G.R. (1997). Cre-mediated chromosome loss in mice. Nature genetics 17, 223-225.

Lionnet, T., Czaplinski, K., Darzacq, X., Shav-Tal, Y., Wells, A.L., Chao, J.A., Park, H.Y., de Turris, V., Lopez-Jones, M., and Singer, R.H. (2011). A transgenic mouse for in vivo detection of endogenous labeled mRNA. Nature methods 8, 165-170.

Maciejowski, J., Li, Y.L., Bosco, N., Campbell, P.J., and de Lange, T. (2015). Chromothripsis and Kataegis Induced by Telomere Crisis. Cell 163, 1641-1654.

Nam, H.S., and Benezra, R. (2009). High levels of Id1 expression define B1 type adult neural stem cells. Cell Stem Cell 5, 515-526.

Nicholson, J.M., Macedo, J.C., Mattingly, A.J., Wangsa, D., Camps, J., Lima, V., Gomes, A.M., Doria, S., Ried, T., Logarinho, E., et al. (2015). Chromosome mis-segregation and cytokinesis failure in trisomic human cells. Elife 4.

Oromendia, A.B., Dodgson, S.E., and Amon, A. (2012). Aneuploidy causes proteotoxic stress in yeast. Genes Dev 26, 2696-2708.

Passerini, V., Ozeri-Galai, E., de Pagter, M.S., Donnelly, N., Schmalbrock, S., Kloosterman, W.P., Kerem, B., and Storchova, Z. (2016). The presence of extra chromosomes leads to genomic instability. Nat Commun 7, 10754.

Pfau, S.J., Silberman, R.E., Knouse, K.A., and Amon, A. (2016). Aneuploidy impairs hematopoietic stem cell fitness and is selected against in regenerating tissues in vivo. Genes Dev 30, 1395-1408.

Schvartzman, J.M., Sotillo, R., and Benezra, R. (2010). Mitotic chromosomal instability and cancer: mouse modelling of the human disease. Nature reviews Cancer 10, 102-115.

Selvarajah, S., Yoshimoto, M., Park, P.C., Maire, G., Paderova, J., Bayani, J., Lim, G., Al-Romaih, K., Squire, J.A., and Zielenska, M. (2006). The breakage-fusion-bridge (BFB) cycle as a mechanism for generating genetic heterogeneity in osteosarcoma. Chromosoma 115, 459-467.

Sheltzer, J.M., Blank, H.M., Pfau, S.J., Tange, Y., George, B.M., Humpton, T.J., Brito, I.L., Hiraoka, Y., Niwa, O., and Amon, A. (2011). Aneuploidy drives genomic instability in yeast. Science 333, 1026-1030.

Sheltzer, J.M., Ko, J.H., Replogle, J.M., Habibe Burgos, N.C., Chung, E.S., Meehl, C.M., Sayles, N.M., Passerini, V., Storchova, Z., and Amon, A. (2017). Single-chromosome Gains Commonly Function as Tumor Suppressors. Cancer Cell 31, 240-255.

Simon, J.E., Bakker, B., and Foijer, F. (2015). CINcere Modelling: What Have Mouse Models for Chromosome Instability Taught Us? Recent Results Cancer Res 200, 39-60.

Sotillo, R., Hernando, E., Diaz-Rodriguez, E., Teruya-Feldstein, J., Cordon-Cardo, C., Lowe, S.W., and Benezra, R. (2007). Mad2 overexpression promotes aneuploidy and tumorigenesis in mice. Cancer Cell 11, 9-23.

Stingele, S., Stoehr, G., Peplowska, K., Cox, J., Mann, M., and Storchova, Z. (2012). Global analysis of genome, transcriptome and proteome reveals the response to aneuploidy in human cells. Molecular systems biology 8, 608.

Tang, Y.C., Williams, B.R., Siegel, J.J., and Amon, A. (2011). Identification of aneuploidy-selective antiproliferation compounds. Cell 144, 499-512.

Torres, E.M., Sokolsky, T., Tucker, C.M., Chan, L.Y., Boselli, M., Dunham, M.J., and Amon, A. (2007). Effects of aneuploidy on cellular physiology and cell division in haploid yeast. Science 317, 916-924.

Williams, B.R., Prabhu, V.R., Hunter, K.E., Glazier, C.M., Whittaker, C.A., Housman, D.E., and Amon, A. (2008). Aneuploidy affects proliferation and spontaneous immortalization in mammalian cells. Science 322, 703-709.

Zhang, M., Cheng, L., Jia, Y., Liu, G., Li, C., Song, S., Bradley, A., and Huang, Y. (2016). Aneuploid embryonic stem cells exhibit impaired differentiation and increased neoplastic potential. Embo J 35, 2285-2300.

Zhu, Y., Kim, Y.M., Li, S., and Zhuang, Y. (2010). Generation and analysis of partially haploid cells with Cre-mediated chromosome deletion in the lymphoid system. The Journal of biological chemistry 285, 26005-26012.

